# Age specific impacts of vegetation functional traits on gastro-intestinal nematode parasite burdens in a large herbivore

**DOI:** 10.1101/2023.01.26.525546

**Authors:** Ellis Wiersma, Robin J. Pakeman, Xavier Bal, Jill G. Pilkington, Josephine M. Pemberton, Daniel H. Nussey, Amy R. Sweeny

## Abstract

1. Gastro-intestinal nematode (GIN) parasites play an important role in the ecological dynamics of many animal populations. Recent studies suggest fine-scale spatial variation in GIN infection dynamics are important in wildlife systems, but the environmental drivers underlying this variation remain poorly understood.
2. We used data from over two decades of GIN parasite egg counts, host space use, and spatial vegetation data from a long-term study of Soay sheep on St Kilda to test how spatial autocorrelation and vegetation in an individual’s home range predict parasite burden across three age groups. We developed a novel approach to quantify the plant functional traits present in a home range to describe the quality of vegetation present.
3. Effects of space and vegetation varied between age classes. In immature lambs, strongyle parasite faecal egg counts (FEC) were spatially structured, being highest in the north and south of our study area. Independent of host body weight and spatial autocorrelation, plant functional traits predicted parasite egg counts. Higher egg counts were associated with more digestible and preferred plant functional traits, suggesting the association could be driven by host density and habitat preference.
4. In contrast, we found no evidence that parasite FEC were related to plant functional traits in the host home range in yearlings or adult sheep. Adult FEC were spatially structured, with highest burdens in the north-east of our study area, while yearling FEC showed no evidence of spatial structuring.
5. Our findings support the importance of fine-scale environmental variation for wildlife disease ecology and provides new evidence that such effects may vary across demographic groups within a population. Parasite burdens in immature individuals appear more readily influenced by fine-scale spatial variation in the environment, highlighting the importance of such heterogeneity for our understanding of wildlife epidemiology and health.

## INTRODUCTION

Gastro-intestinal nematode (GIN) parasites play an important role in the ecology of wildlife populations, harming the host, and impacting individual fitness and population dynamics (Coulson *et al*., 2018). Studies in livestock systems provide detailed understanding of the factors influencing GIN transmission and the development and maintenance of immunity under relatively controlled environmental conditions (Roeber *et al*., 2013; Smith *et al*., 2009; Vlassoff *et al*., 2001), but without incorporating the complexity evident in wild systems. Host factors such as age, sex, and body condition are well-established predictors of GIN burdens in wildlife systems (Lynsdale *et al*., 2017). However, recent studies have identified complex fine-scale patterns of spatial and temporal variation in parasite burden over and above the effects of these host variables (Albery *et al*., 2019; Albery *et al*., 2020; Sweeny, Albery *et al*., 2021; Verheyden *et al*., 2020). While variation in host habitat use is one potential explanation for spatial variation in parasite burdens in the wild (e.g., Carbayo *et al*., 2019), our understanding of the factors regulating variation in GIN parasite burdens in natural populations experiencing complex spatially and temporally variable environmental conditions remains limited, representing an important gap in our understanding of wildlife disease epidemiology.

Transmission of GIN is driven both by host factors influencing susceptibility and by environmental factors influencing exposure. Most GIN have a lifecycle involving adult males and females living and mating in host guts, producing eggs that are shed via the faeces into the host’s environment. These eggs develop into free-living larval stages, which are ingested by new hosts and then develop into adults (Morand *et al*., 2007). Both the rate of egg shedding by adult females and the ingestion of infectious larvae by susceptible hosts via contaminated food or water work in tandem to drive helminth dynamics (Brooker *et al*., 2006; Morand *et al*., 2007). Furthermore, environmental conditions can influence infectious life stages of GIN outside the host, as well as resource availability and host nutritional state, both of which may be altered over space and time (Becker *et al*. 2020). Incorporating measures of host and parasite habitat can offer additional insights into complex exposure and susceptibility drivers of host-parasite dynamics varying over spatial and temporal scales.

Herbivore grazing strategies and the faecal-oral transmission of GIN makes herbivores especially vulnerable to spatial variation in fine-scale habitat and vegetation quality effects on GIN risk and burden. Vegetation type impacts the decomposition of herbivore faeces (Williams & Warren, 2004), and GIN eggs are sensitive to decomposition rates (Nielsen *et al*., 2007), suggesting that the structure and type of vegetation can directly impact GIN survival and hence, generate strong spatial structure of transmission risk. For example, taller grass swards can provide more stable microclimates for GIN eggs and larvae (Hutchings *et al*., 2002), potentially retaining moisture and reducing rates of faecal breakdown. Indirectly, vegetation quality may shape host condition, immunity status, and ability to resist infection (Budischak *et al*., 2018) or avoid exposure (Hutchings *et al*. 2001). For example, female red deer with higher quality home ranges show higher survival probability (Froy *et al*. 2018), and wild bovids with lower-quality diets have more GI parasites during drought years (Ezenwa, 2004). However, areas of highest quality grazing may be associated with higher transmission risk, as high-quality habitats are likely subjected to increased competition for forage and higher densities of individual grazers. This could counter any beneficial effects on host condition associated with high quality vegetation and increase transmission due to high levels of GIN exposure. While recent field studies of GIN infection dynamics have detected and accounted for spatial autocorrelation in parasite burden (e.g., red deer, Albery *et al*., 2018), our understanding of the environmental drivers of spatial variation is limited. Here, we take a novel functional trait-based approach to characterise the vegetation within the home ranges of individuals to test the extent to which fine-scale variation in host habitat may shape GIN dynamics.

Describing herbivore habitats in terms of plant species’ functional traits – their morphological, physiological, and phenological features – can quantify plant species’ contribution to ecosystem processes (Roscher *et al*., 2012; Tilman, 2001) and offer a useful way of making broad ecological predictions without reference to specific species. The mass-ratio hypothesis states that the dominance and prevalence of certain traits largely determines an ecosystem’s functioning (Díaz *et al*., 2007; Grime, 1998); effectively, the contribution of a species to ecosystem function depends on its trait values and its contribution to biomass in that trophic level. Using a trait-based approach allows variation in species’ identities between systems to be standardized (Weiss & Ray, 2019) and provides a predictive framework within which to work (McGill *et al*., 2006). For example, both specific leaf area (SLA) (Garnier *et al*. 2004) and leaf dry matter content (LDMC) (Pakeman, 2014) have been used as a proxy for productivity. Quantifying the functional traits of plants present within a host’s home range can improve understanding of how habitat and food quality influences parasite burdens in wildlife.

The Soay sheep (*Ovis aries* L.) of St. Kilda are a valuable study system to test how habitat quality influences parasite burdens, due to long-term data on individual space use, GIN burden, and morphometric measurements (Wilson *et al*., 2004). These sheep are hosts to a range of GIN parasites, predominantly *Teladorsagia circumcincta, Trichostrongylus axei* and *Trichostrongylus vitrinus* (Craig *et al*., 2006), and parasite burdens can be estimated using standard faecal egg counting (FEC) techniques (Cabaret *et al*., 1998; Cringoli *et al*., 2009). Previous studies of the Soay sheep have established that heavy GIN burdens are associated with gastro-intestinal damage and higher over-winter mortality risk (Gulland, 1992) and with negative fitness consequences in lambs (Hayward *et al*., 2011). FEC varies with age: it is highest in immunologically naïve lambs, declines and stabilises at prime age (2-5 years) as immunity to GIN develops, and then increases again in later adulthood (6 years onwards) as immunity wanes (Hayward *et al*., 2009). Males tend to have higher burdens than females (Wilson *et al*., 2004). Recently, a detailed map of the spatial distribution of plant species within our study area found that the proportion of *Holcus lanatus* – a common, comparatively high quality, easily digestible grass species – cover in an individual’s home range predicted lifetime breeding success (Regan *et al*., 2016).

In this study, we aimed to determine if the vegetation functional traits within an individual’s home range were predictive of variation in GIN burdens (measured as FEC) and whether associations are age dependent. We controlled for host weight and spatial patterns of parasite infection to detect effects of vegetation above and beyond host position in space and to rule out indirect effects on condition. We predicted that individuals whose home range contained more digestible and nutritive leaves would have higher GIN burdens as they are associated with more preferred vegetation and hence higher faecal inputs. We also expected taller vegetation, greater leaf size, and species indicative of high moisture conditions to have higher FEC through enhanced parasite survival in their free-living stages. As lambs tend to have little acquired immunity, we expected lambs to be more sensitive to vegetation traits and environmental influences than adults.

## METHODS

### Study system

The Soay sheep of the St. Kilda archipelago (57°49′N, 08°34′W, 65 km NW of the Outer Hebrides, Scotland; Figure 1) are an isolated and unmanaged population (Clutton-Brock *et al*., 2004). Individuals in the Village Bay area of the largest island, Hirta, have been individually marked and monitored since 1985 with over 95% of individuals in this area being individually marked at any time (Clutton-Brock *et al*., 1992). Each year, lambs are caught shortly after birth in April (typically within a week), marked with unique ear tags and weighed. In August, as many individuals as possible are caught in corral traps over a two-week period. Typically, 50-60% of the resident Village Bay sheep population is captured each August (Clutton-Brock & Pemberton, 2004) and morphometric measurements are taken, including body mass (to the nearest 100 g). August body mass is an individually repeatable and heritable trait, which is negatively associated with August FEC and positively associated with survival in the following winter (Clutton-Brock *et al*., 1992; Craig *et al*., 2008). All fieldwork was conducted in accordance with the Animals (Scientific Procedures) Act 1986 and with permission from the University of Edinburgh Animal Welfare Ethical Review Body. All sampling was carried out in accordance with UK Home Office regulations under Project License PP4825594.

**Figure 1.**
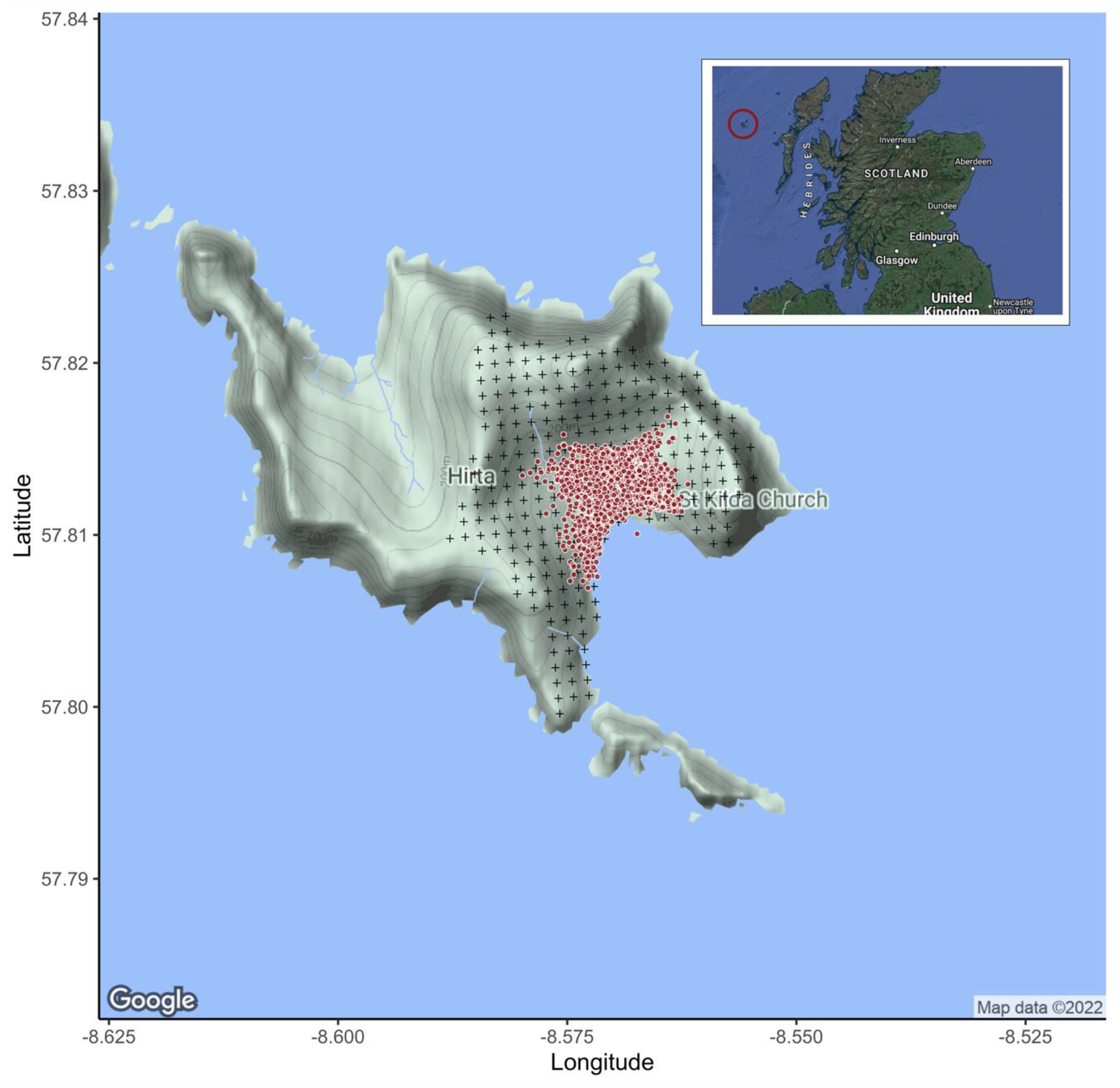
Map of St. Kilda with Village Bay in the southeast region of the island. Hectares used in vegetation surveys and censusing marked by grid. Dots indicate average census location of individual sheep as a relative representation of sheep home ranges.

Faecal samples are collected rectally at capture in August or, where a rectal sample could not be obtained, from observed defecation within several days of capture. As GIN eggs cannot be distinguished to species by eye, eggs from *Teladorsagia* spp., *Trichostrongylus* spp., *Chabertia ovina, Bunostomum trigonocephalum, and Strongyloides papillosus* (hereafter referred to as strongyles) are combined and counted as one FEC using a modification of the McMaster technique (Wilson *et al*., 2004). Actual strongyle burden and FEC are strongly correlated, with lambs having higher parasite burdens than adults (Wilson *et al*., 2004). We used August FEC data collected from 1988-2017 from lambs (Males: N=870; Females: N=891), yearlings (Males: N=356; Females: N=510), and adults (Males: N=547; Females: N=2336) where associated August weight and annual home range data was available. Sample sizes indicated here are unique individual-year samples. For individuals with multiple FEC data points for the same year, we kept the earliest dated sample and removed all others. We also removed FEC values greater than 5000 to minimise overdispersion (3 data points removed) resulting in 1081 individual adults with 2883 total observations, 866 yearlings, and 1761 lambs. Unbalanced sample sizes between sexes and ages are due to higher mortality in males and younger individuals resulting in fewer samples, as well as greater dispersal in males (Clutton-Brock *et al*., 2004).

### Vegetation and plant functional traits

The study area contains approximately 30% of the island’s total sheep population (Clutton-Brock *et al*., 1992). The vegetation of the study area is dominated by *Holcus lanatus*–*Agrostis capillaris* (*HA*) grassland on free draining, fertile soils, particularly around the village, *Molinia caerulea*-dominated grassland on areas receiving drainage, and *Calluna vulgaris-*dominated heath (wet and dry) on the steeper slopes around the bay (Jewell *et al*. 1974; Vicari *et al*., 2018) (Supplementary Figure S1). HA grassland has the highest live standing-crop biomass of the area (Crawley *et al*. 2004; Crawley *et al*., 2021) and is preferred by Soay sheep (Jones *et al*., 2006).

Wet *Calluna* heathland is rarely grazed even in years of resource limitation due to high sheep population numbers (Crawley *et al*., 2021). Vegetation surveys were conducted from 2008–2012 by visually estimating the proportion of species cover for every hectare (N=267) to the nearest 5% with all estimates done by one observer. Species composition of the vegetation community remained relatively constant between 1993 and 2012 (summarised in Crawley, 2017).

We chose four plant functional traits and two measures of plant habitat preferences to characterise the vegetation (Table 1). These traits were chosen for their potential relevance to parasite transmission or disease susceptibility, for example, due to increased digestibility of vegetation for grazers or more suitable microclimates for strongyle larvae. LDMC, SLA, and Ellenberg nitrogen content (N) can all be used as proxies for preference/digestibility, as sheep tend to favour vegetation which are lacking fibrous compounds, easy to digest, and with high nutritional content (Gardarin *et al*., 2014; Pakeman, 2014). Canopy height (CanHt), Ellenberg moisture value (F), and leaf size (LeafSize) could be associated with enhancing parasite survival in their free-living stages with high values for each indicating favourable microclimate conditions (Table 1). Trait values for each species were obtained from UK calibrated Ellenberg values (F and N) (Hill *et al*. 1999) and the LEDA Traitbase (CanHt, SLA, LeafSize, and LDMC) (Kleyer *et al*., 2008). Community Weighted Means (CWM) – an average trait value of an area – were calculated for each hectare of the study area based on the functional trait values of each species and weighted by the area each species occupied using the *FD* package v.1.0-12 (Laliberté and Legendre, 2010; Laliberté *et al*., 2014). The range of CWMs for CanHt, LeafSize, and F were spread across the study area. Low LDMC, high SLA, and high N were concentrated around the former settlement in the Village Bay area (Supplementary Figure S1). To account for correlations between functional traits (Supplementary Table S1), we ran a principal component analysis using the *stats* package within R (R Core Team, 2021) to generate a combined measure of grazing preference. LDMC, LeafSize, N, and SLA traits were collapsed into a principal component (PC1) and carried forward for further analysis (see Results).

**Table 1.** Definitions and predicted effects of plant functional traits used.

| Term | Description | Predicted effect on GIN transmission/survival | Predicted effect on sheep condition |
| --- | --- | --- | --- |
| CanHt | Distance between the base of a plant and its highest photosynthetic tissue (m). <sup>1</sup> | Positive: more favourable microclimate conditions | Positive: greater nutrition for given bite size |
| LDMC | Ratio of dry mass to fresh mass (mg/g). <sup>2</sup> | Negative: less palatable vegetation reducing host density | Negative: reduced palatability at high values, greater nutrition at low values |
| Leaf Size | Area of one side of a fresh leaf (mm<sup>2</sup>). <sup>1</sup> | Positive: more favourable microclimate conditions | Positive: greater nutrition for given bite size |
| SLA | Area of one side of a fresh leaf divided by the dry weight (mm<sup>2</sup>/mg). <sup>1</sup> | Positive: more palatable vegetation increasing host density | Positive: increased palatability at high values |

<sup>1</sup> Weiher *et al.*, 1999; <sup>2</sup> Wilson *et al.*, 1999; <sup>3</sup> Hill *et al.*, 1999

| Term | Description | Predicted effect on GIN transmission/survival | Predicted effect on sheep condition |
| --- | --- | --- | --- |
| F | Ellenberg moisture indicator (on a scale of 1 to 9). Low values indicate very low soil moisture content.<sup>3</sup> | Positive: more favourable microclimate conditions | - |
| N | Ellenberg nitrogen indicator (on a scale of 1 to 9). Low values indicate highly infertile sites.<sup>3</sup> | Positive: more palatable vegetation increasing host density | Positive: greater nutrition |

<sup>1</sup> Weiher *et al.*, 1999; <sup>2</sup> Wilson *et al.*, 1999; <sup>3</sup> Hill *et al.*, 1999

### Home range quality estimation

30 study area censuses are carried out each year (ten each in spring, summer, and autumn). During each census, all individuals seen along a set route are noted, along with their location to the nearest 100 m x 100 m Ordnance Survey grid square. Adult Soay sheep tend to segregate into social groups according to sex, with females exhibiting greater philopatry to their birth area than males. While male juveniles leave their mothers at approximately six months old, females remain in the vicinity of their mother until about two years old (Clutton-Brock *et al*., 2004).

We calculated home ranges based on census sightings from 1988–2017; years for which FEC data was available. We used the R package *adehabitatHR* v.0.4.19 (Calenge, 2006) to estimate home ranges. Coordinates for census points are recorded as the most south-westerly corner of a pre-determined hectare, so we corrected the coordinates to give the centre of each hectare (easting + 50 m, northing + 50 m). As many observations have identical hectare references, we added a random number between –20 and 20 (a jitter of up to 20 m) to both easting and northing coordinates of each census observation to prevent errors with kernel estimates (Regan *et al*., 2016). To maximise home range accuracy, we used all observations for an individual with at least 10 census observations in a year (Number of individuals per year with home range and parasite data: mean: 183.7; standard deviation: 64.4; range: 28–294). We used a 70% isopleth to calculate the core home range (Powell, 2000; Regan *et al*., 2016). We then overlaid the core home range onto the vegetation hectare squares previously developed. To account for different contributions from each hectare square to home ranges we used proportional weighting of the number of observations of every individual in each square. This resulted in a mean value for each functional trait for each individual sheep’s home range.

### Statistical analysis

We conducted all statistical analyses with August FEC as our response variable in R v.4.0.4 (R Core Team, 2021). Individuals were categorised as adults, yearlings, or lambs due to the pronounced variation in FEC across the age groups (Sweeny *et al*., 2022). Spatial autocorrelation, where samples or individuals closer together in time and space are more alike than those further apart, is becoming increasingly important to consider in studies of disease ecology (Albery *et al*., 2022). To quantify spatial autocorrelation, we included a stochastic partial differentiation equation (SPDE) random effect in Integrated Nested Laplace Approximation (INLA) models from the *INLA* package v.21.02.23 (Lindgren *et al*., 2011; Rue *et al*., 2009). To do so, we first developed a separate spatial mesh for adults, yearlings, and lambs constrained by the coastline boundaries. Each age-specific mesh is made up of the average coordinate for each individual-year home range and allows interpolation of spatially distinct samples. Any average coordinate that fell in the ocean was removed (N=6). All subsequent models were run using the age-appropriate mesh.

The base model for all age categories included fixed effects of sex and August weight, and year as a random effect. SPDE (spatial field) was added to models separately as a random effect to test the effect of spatial autocorrelation. The model for the adult age group also included age as a continuous covariate and individual identity as a random effect to account for pseudo replication. All models were fitted with negative binomial distributions. To base + SPDE models for each age group, we added vegetation traits – PC1, CanHt, and F – separately. If a trait improved the model fit, we then added all combinations of pairs of traits to the model to test whether effects were independent of one another. We compared the fit of models using DIC and identified the best fitting model as the one with the fewest parameters within 2 DIC units of the model with lowest overall DIC.

## RESULTS

### Principal component analysis

Analysis of correlations among the vegetation functional trait scores of each of the study area grid squares identified strong correlations among N, SLA, LeafSize and LMDC (Supplementary Table S1). Principal component analysis confirmed this and identified a first principal axis of variation (PC1) explaining 57.1% of trait variance (Figure 2) with strong loadings in one direction from LMDC and in the opposite direction from LeafSize, SLA and N (Figure 2, Supplementary Table S2). PC1 is consistent with expected sheep dietary preferences, with high values of PC1 indicating tougher to digest and less nutritious vegetation, and we chose to include it as a single variable in subsequent analysis. CanHt and F both had higher loadings on PC2 than PC1, but they were almost orthogonal to each other, and so were used in the model as separate terms rather than using the aggregated PC2. F also had a relatively high loading on PC1 and was correlated to the variable associated with PC1, indicating some confounding between F and PC1 was possible; wetter areas were also characterised by tougher, harder to digest leaf material.

**Figure 2.**
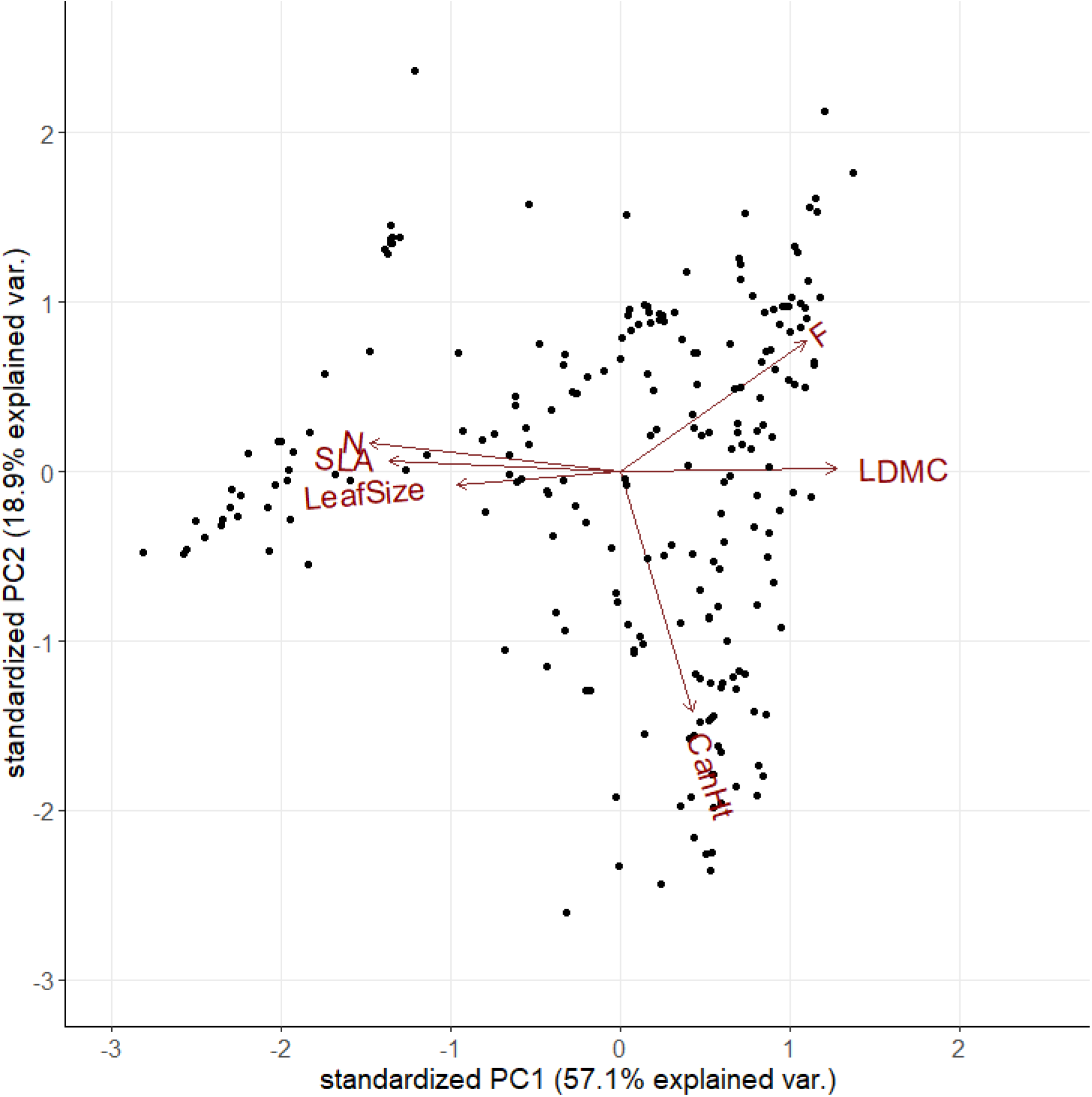
Principal component analysis (PCA) of functional trait values calculated for each hectare of the study area. The total variance explained by the first two axes is 76.0%. The loading scores and contributions of each variable are given in Supplementary Table S2.

### FEC Models and effect sizes

We found evidence for effects of spatial autocorrelation in August FEC for lambs and adults, but not for yearlings (δDIC: adults: 2.53; yearlings: −1.18; lambs; 4.27; Table 2). In lambs, we found additional evidence for vegetation trait effects beyond spatial structuring of FEC (Table 2, Figure 3). In contrast, the best supported model in terms of DIC for yearlings and adults did not include any of the vegetation traits (adults: Base + SPDE; yearlings: Base; Table 2). As expected, males had higher FEC than females and weight was negatively associated with FEC across age groups, although there was no effect of age in adults (Figure 3, Supplementary Figure S2). For lambs, inclusion of the spatial effect improved model fit (δDIC: 4.27, Table 2) and the addition of either F or PC1 to the SPDE model provided further improvement (δDIC: F: 13.03; PC1: 13.63, Table 2). Estimates from these models revealed significant negative effects of both F and PC1 when included on their own in the FEC models (Figures 3 and 4). However, the models including F or PC1 explained very similar levels of variance in FEC (δDIC: 0.6, Table 2) and including them together in the model slightly increased the DIC (Table 2). This suggests the two vegetation scores were explaining largely shared variance in FEC. Examination of the spatial field of the lamb models including F or PC1, which would illustrate spatial autocorrelation in FEC after accounting for effects of the vegetation traits, weight, and sex, suggest higher strongyle densities in the north and south of the study area (Figure 5A,B). Adult spatial fields suggest high strongyle densities in the north-east of the study area (Figure 5C).

**Table 2.** DIC values of model construction. Models are presented in sequential order of term addition. Base models include sex, weight, and year for all age categories, and age and individual ID for adults only. Terms were retained in the model if DIC values decreased by >2 δDIC. Final models (indicated in bold) were chosen based on DIC values (>2 δDIC) and least term parsimony.

| Age Category | Model | DIC | $\delta DIC$ |
| --- | --- | --- | --- |
| Adult | Base | 27611.71 | 0.00 |
|  | **Base+SPDE** | **27609.18** | **2.53** |
|  | Base+SPDE+CanHt | 27612.07 | -0.36 |
|  | Base+SPDE+F | 27611.74 | -0.03 |
|  | Base+SPDE+PC1 | 27613.63 | -1.92 |
| Yearling | **Base** | **11245.89** | **0.00** |
|  | Base+SPDE | 11247.07 | -1.18 |
|  | Base+SPDE+CanHt | 11249.06 | -3.17 |
|  | Base+SPDE+F | 11249.92 | -4.03 |
|  | Base+SPDE+PC1 | 11249.34 | -3.45 |
| Lamb | Base | 26315.04 | 0.00 |
|  | Base+SPDE | 26310.77 | 4.27 |
|  | Base+SPDE+CanHt | 26309.92 | 5.12 |
|  | **Base+SPDE+F** | **26302.01** | **13.03** |
|  | **Base+SPDE+PC1** | **26301.41** | **13.63** |
|  | Base+SPDE+CanHt+F | 26303.36 | 11.68 |
|  | Base+SPDE+CanHt+PC1 | 26303.35 | 11.69 |
|  | Base+SPDE+F+PC1 | 26302.93 | 12.11 |

**Figure 3.**
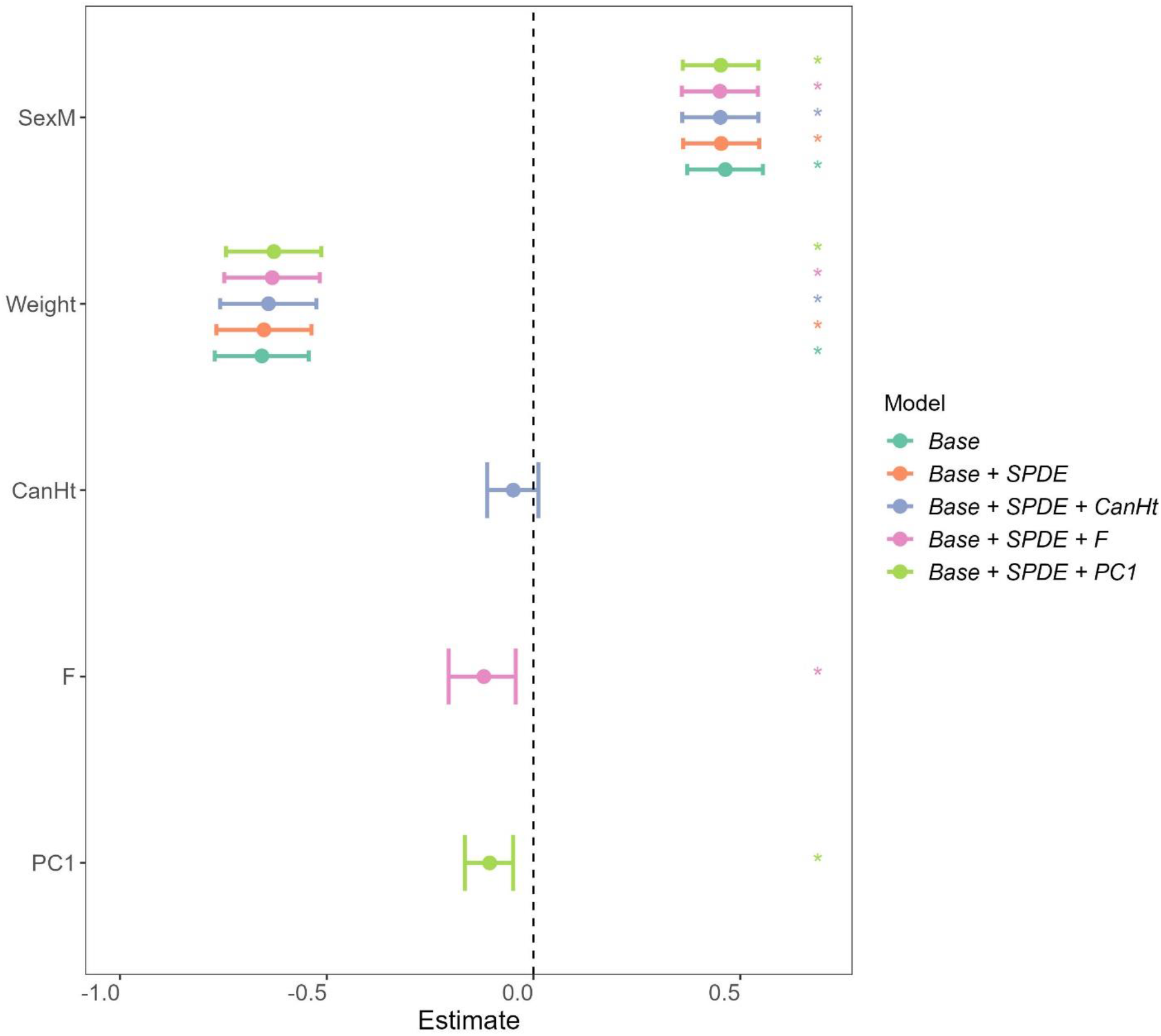
Confidence intervals (95%) for models on lambs. Base model includes sex, and weight as fixed effects, and year as a random effect. The similarity in confidence intervals of each term across models suggests that the effects are consistent following inclusion of random effects. Each model, and the combination of terms included therein, is assigned a colour and, as not all models include all terms, not all models can be compared for a given term. Traits are insignificant in the model if the confidence interval crosses the central dashed line. Positive estimates indicate a positive effect of the term on strongyle burdens.

**Figure 4.**
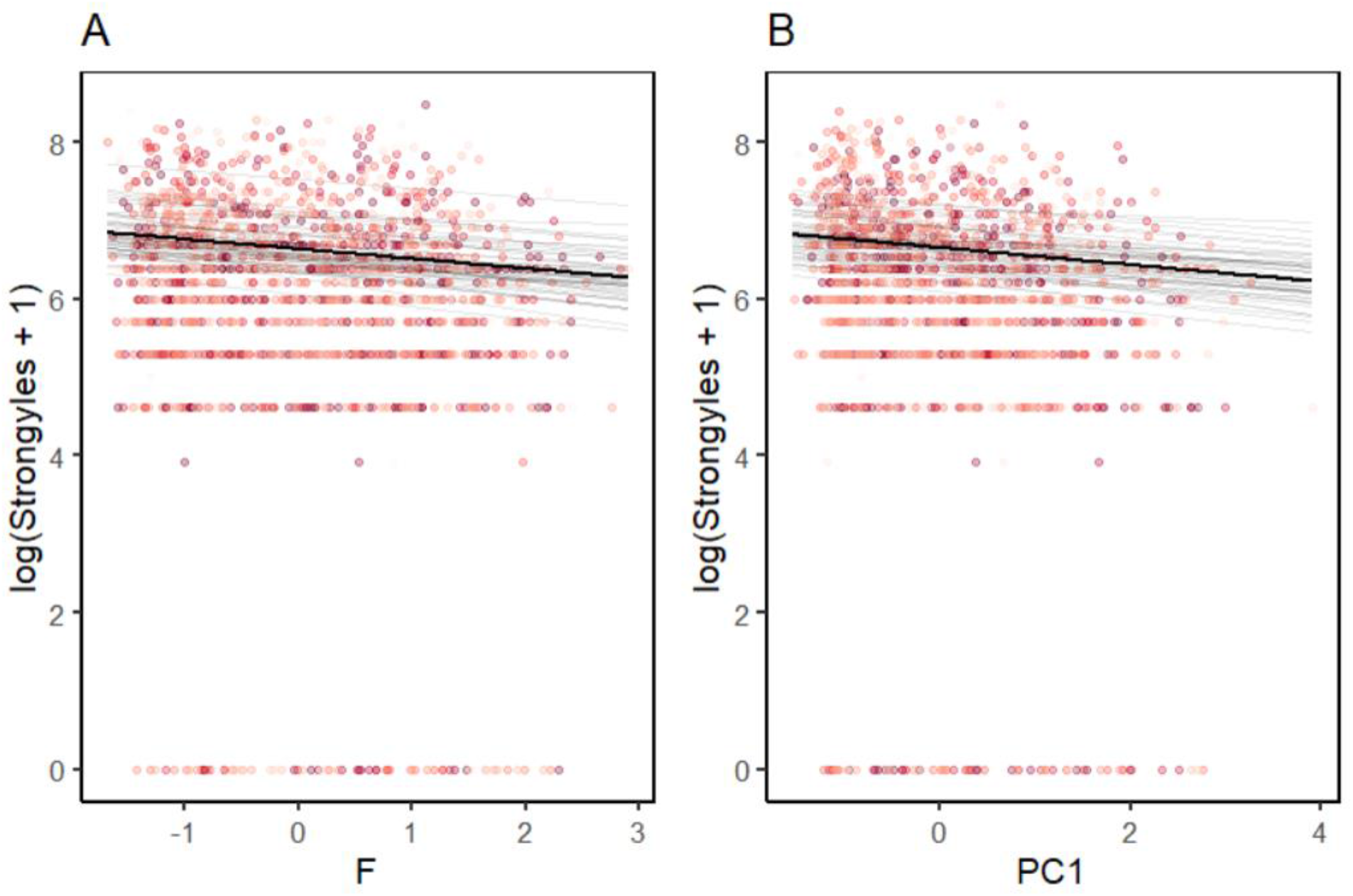
Effect of vegetation functional traits (A) F and (B) PC1 on FEC in lambs from final model outputs. The functional trait values are standardized around 0 and show a significant decrease in strongyle burdens as trait values increased. F: Mean: −0.125; 95% confidence interval: (−0.213, − 0.047); p=0.00124. PC1: Mean: −0.11; 95% confidence interval: (−0.171, −0.052); p=0.00023.

**Figure 5.**
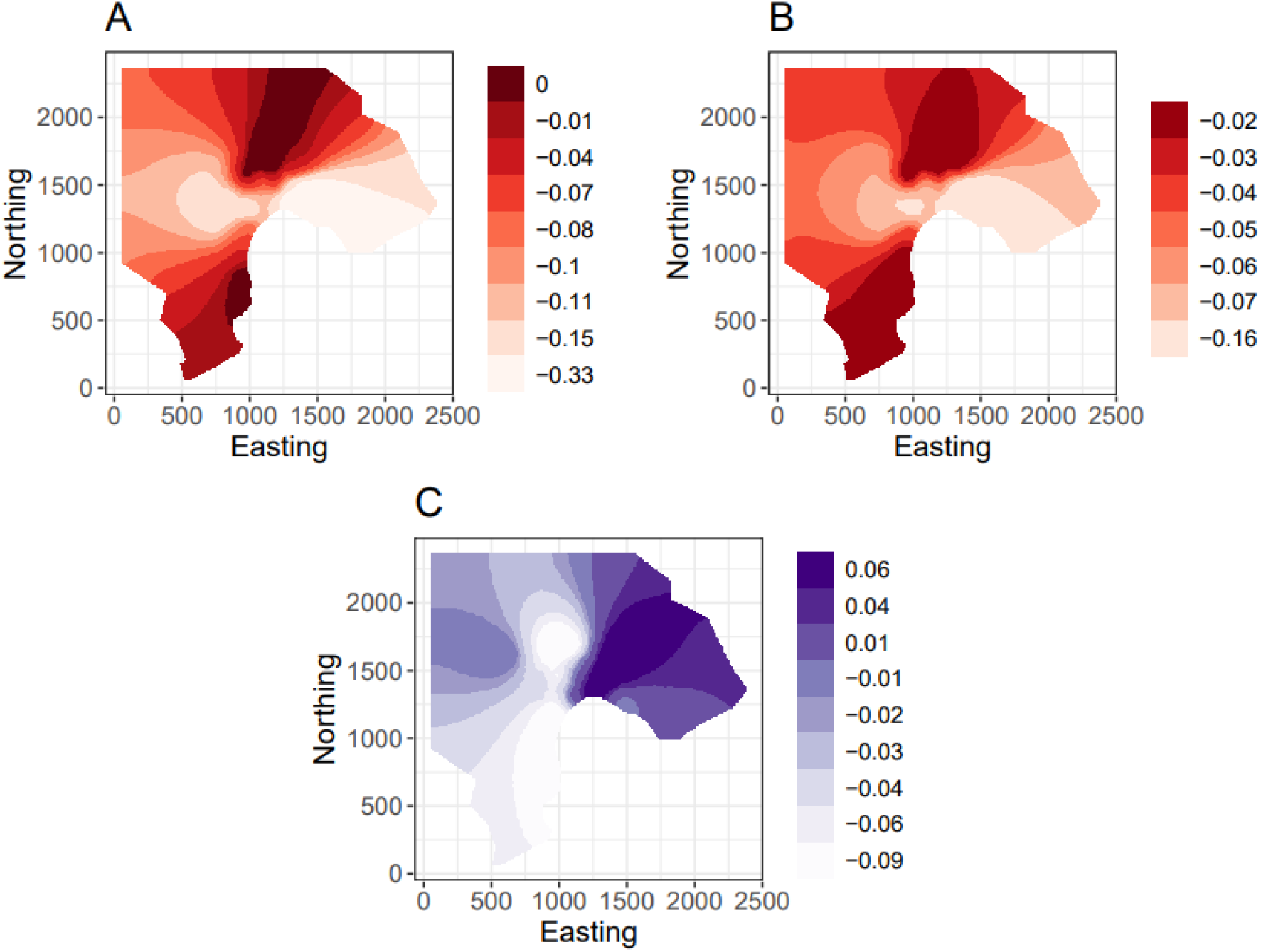
Spatial fields for lamb and adult FEC models. Lamb spatial fields models include base terms and functional traits (A) F or (B) PC1. Adult spatial field model (C) includes base terms. Colour scale shows the lower bounds of the spatial effect quantiles with darker colours indicating higher parasite counts. Eastings and northings are standardised coordinates (in units of 1 m) from the southern- and western-most points of the study area and the spatial fields cover the Village Bay area (see Figure 1).

## DISCUSSION

We found age-dependent effects of both vegetation traits and spatial autocorrelation on strongyle burdens in Soay sheep. In lambs, vegetation traits (F and PC1) were negatively associated with strongyle burdens. Strongyle burdens in adults and yearlings showed no association with vegetation. Yearlings were also the only age category where accounting for spatial autocorrelation did not significantly improve model fit, with high parasite burdens in the north-east, and in the north and south of the Village Bay area for adults and lambs respectively. Our findings confirm the importance of fine-scale spatial variation in the landscape ecology of disease in natural systems and offer a new approach to understanding the relationship between habitat and infection status for field studies of herbivores. Importantly, we show that spatial variation and effects of habitat on host-parasite systems can vary among host demographic groups and highlight the importance of understanding this among-host variation for disease ecology.

### Parasite associations with PC1 and F in lambs

High values of PC1 (high LDMC and low N, SLA, and LeafSize) reflect low preference vegetation with low digestibility. The effects of PC1 on lamb strongyle burdens were as expected: higher strongyle burdens were associated with more digestible vegetation traits. It is unlikely that this is a condition-mediated effect as we included weight in our models – weight has consistently been shown as a good predictor of individual condition in this study system (Jones *et al*., 2005; Milner *et al*., 1999) – and areas of good grazing should then be associated with more resources for immunity and lower FEC. This suggests that exposure to infective larval stages could be more important in determining parasite burdens than individual condition. While experimental work in wood mice (*Apodemus sylvaticus*) showed increased resistance to helminth infection following diet supplementation (Sweeny, Clerc *et al*., 2021), nutrition effects can operate through both exposure and susceptibility processes and produce various outcomes for disease prevalence (Becker & Hall, 2014). In Soay sheep, much of the variation in parasite burdens is likely down to landscape heterogeneity with sheep density driving greater parasite exposure and transmission between individuals. However, since we controlled for spatial autocorrelation in our models, it is likely that there are further mechanisms behind transmission than solely spatial location.

One theory could be that social grouping and dynamics play a larger role than spatial location. Soay sheep do show some sex-based social segregation; males tend to form a dominance hierarchy, and male lambs tend to leave their mothers earlier than female lambs (Clutton-Brock *et al*., 2004). Additionally, adult males have a different seasonal FEC pattern to lambs or adult females, potentially leading to altered transmission and infection dynamics (Sweeny *et al*., 2022). Incorporating social networks and accounting for how different interactions (e.g., mother-offspring relationships) may drive disease transmission could provide further information behind the mechanisms of such transmission. Previous work on the European badger (*Meles meles*) provides a framework for such work by comparing spatial models with social interaction models to determine which best described parasite distribution (Albery *et al*., 2020).

We found that high levels of moisture (F) were associated with lower parasite burdens. While contrary to our expectations – rainfall has been found to favour strongyle larval survival (Zouyed *et al*., 2018) – moisture may not be limiting strongyle survival on St. Kilda due to its hyperoceanic climate. Instead, sheep may avoid the wettest areas (Putfarken *et al*., 2008), thus reducing rates of host density-dependent transmission and leading to a negative association between moisture levels and strongyle burdens. There may also be some unknown influence of fine-scale spatial heterogeneity of the plant community on parasite burdens (Johnson *et al*., 2016). We used averaged vegetation traits over the entirety of an individual’s home range and so can only provide a limited picture of the heterogeneity of vegetation available to an individual. Sheep are selective about their food sources (Illius *et al*., 1995), so simply because their home range has a particular trait value does not mean the plants contributing to this will be consumed. Additionally, greater spatial heterogeneity tends to increase community stability in both theoretical models (Briggs & Hoopes, 2004; Murdoch, 1977), empirical predator-prey systems (Daugherty, 2011; Mitsunaga & Fujii, 1997), and host–parasite interactions (Brockhurst *et al*., 2006). Therefore, small-scale structural and environmental variation in habitat must be considered along with large-scale generalisations of habitat quality in studies of wildlife disease and ecology.

### Age-specific vegetation-parasite associations

We found support for spatial structuring of FEC in lambs and adults, and effects of vegetation traits on parasite counts only in lambs, but little support for either in yearlings. To our knowledge, this is the first evidence that fine-scale habitat effects on parasite burdens might vary across demographic groups in the wild. This age-specific effect of space and vegetation on strongyle burdens may be driven by differences in foraging behaviour between age classes. Young, domesticated cattle learn to graze faster when in the presence of experienced individuals (Costa *et al*., 2016), and moose (*Alces alces*) calves learn to identify toxic plants from their mother (Edwards, 1976). Additionally, as the vegetation trait values used in this study are average values and do not consider intraspecies variation, it may be that adults are more adept at identifying the variation that minimises any nutrition-parasite trade-off. For example, Soay sheep preferentially graze shorter plants with a comparatively reduced parasite risk, over taller plants associated with greater nutritional intake (Hutchings *et al*., 2002). As lambs may not yet have developed learned behaviours regarding grazing, this suggests that they may not have learnt to avoid vegetation with higher parasite transmission risk. Therefore, their parasite burdens may be more correlated with functional traits at the scale of this investigation than adults, who are more selective consumers due to these learned behaviours. As lambs are dependent on their mother and tend follow her, there could be a potential conflict between adult survival and performance from feeding on the best vegetation, and exposure of their offspring to higher parasite burdens. Indeed, Soay sheep lambs are less selective in consuming vegetation and are less avoidant of areas of high larvae count than adults (Hutchings *et al*., 2002).

Lambs are also immunologically naïve and less resilient to infection than adults (Nussey *et al*., 2012). Soay sheep rapidly acquire resistance to GINs, with yearlings having higher antibody levels and lower FEC than lambs (Sparks *et al*., 2018). As yearlings are less susceptible to disease than lambs, this may contribute to the lack of spatial structure in yearling FEC. It is also likely that the yearling age group has less variation in individual condition as weak lambs will be less likely to survive their first winter and mature into yearlings, and yearlings are not affected by immunosenescence as in adults. This suggests that strongyle burdens in older individuals are largely driven by variation in immune-mediated resistance or tolerance and condition, while spatial variation of lamb burdens may be more directly linked to eggs ingested from pasture and so more easily discerned. Therefore, including measures of immune response in Soay sheep may provide further insight into whether it is a lack of immune response in lambs or other factors driving this difference between age classes.

### Wider implications and conclusions

We present a novel approach for analysing the interaction between individuals and their environment influences disease dynamics. While incorporating the vegetation of an individual’s home range into analyses of fitness traits has been done previously in the Soay sheep (Regan *et al*., 2016), using vegetation functional traits to summarise an individual’s home range allows for more generalisable conclusions about processes across study systems. In addition, there is growing awareness that fine-scale spatial variation is an important driver of parasite dynamics in natural systems (Albery *et al*., 2021; Becker *et al*., 2020), to which our findings lend further support. Unfortunately, we cannot separate effects of exposure linked to sheep density from effects of exposure linked to vegetation quality using this approach. The effect of PC1 is consistent with high quality grazing areas being more heavily used by sheep and thus higher strongyle burdens in lambs. This means that patterns in strongyle burdens are related to selective grazing patterns and transmission. However, we cannot exclude that this effect may be due to more specific characteristics of the vegetation itself and how this impacts larval development and transmission.

In conclusion, we present fine-scale patterns of variation in parasite burden, which are relatively rarely explored in wild systems (Albery *et al*., 2022). Our models suggest there is both spatial structuring and effects of vegetation type on strongyle burdens which may well be driven by preferential grazing and local density. However, we also show that these patterns are not present in all members of population and may depend on various demographic factors such as age, immunological susceptibility, or condition. This has implications for our understanding of wildlife epidemiology and health as not all individuals or demographic groups experience the same disease risk. In addition, identifying spatial hot spots for parasite transmission could lead to more effective grazing management plans to reduce disease-related costs and dependence on anthelmintics within livestock systems.

## Acknowledgements

We acknowledge the National Trust for Scotland for permission to work on St. Kilda and QinetiQ, Eurest and Kilda Cruises for providing logistics and support for the long-term study. We also thank Ian Stevenson of Sunadal Data Solutions for his tireless and brilliant work building and maintaining our project database, Greg Albery for constructive comments on analyses, and Mike Evans and Mariecia Fraser for comments on manuscript drafts. Mick Crawley planned the hectare-scale botanical recording in 2007 and carried out all field work associated with estimating the percentage cover of each species each August between 2008 and 2012. The long-term study was funded largely by consecutive responsive mode grants from NERC, as well as grants from BBSRC, Wellcome Trust, the Leverhulme trust and ERC. This particular project was enabled by a large NERC grant (NE/R016801/1). We are immensely grateful to the many fieldworkers who have collected data and samples on St Kilda over the years.

## Author Contributions

EW, ARS, DHN and RJP conceived of and designed the study. EW and ARS conducted the analyses. EW, ARS, DHN and RJP prepared the first draft of the manuscript, and all authors contributed to revised drafts. JGP, XB and JMP led the fieldwork on St Kilda and JGP collected most of the faecal egg count data.

## Data Availability Statement

Data will be available from the Dryad Digital Repository following final manuscript acceptance.

## REFERENCES

Albery, G. F., Newman, C., Ross, J. B., MacDonald, D. W., Bansal, S., & Buesching, C. (2020). Negative density-dependent parasitism in a group-living carnivore. Proceedings of the Royal Society B, 287(1941), 20202655. https://doi.org/10.1098/rspb.2020.2655

Albery, G.F., Becker, D.J., Kenyon, F., Nussey, D.H. & Pemberton, J.M. (2019). The fine-scale landscape of immunity and parasitism in a wild ungulate population. Integrative and Comparative Biology, 59(5), 1165–1175. https://doi.org/10.1093/icb/icz016

Albery, G.F., Kenyon, F., Morris, A., Morris, S., Nussey, D.H. & Pemberton, J.M. (2018). Seasonality of helminth infection in wild red deer varies between individuals and between parasite taxa. Parasitology, 145(11), 1410–1420. https://doi.org/10.1017/S0031182018000185

Albery, G.F., Kirkpatrick, L., Firth, J.A. & Bansal, S. (2021). Unifying spatial and social network analysis in disease ecology. Journal of Animal Ecology, 90(1), 45–61. https://doi.org/10.1111/1365-2656.13356

Albery, G.F., Sweeny, A.R., Becker, D.J. & Bansal, S. (2022). Fine-scale spatial patterns of wildlife disease are common and understudied. Functional Ecology, 36(1), 214–225. https://doi.org/10.1111/1365-2435.13942

Becker, D.J., Albery, G.F., Kessler, M.K., Lunn, T.J., Falvo, C.A., Czirják, G.Á., Martin, L.B. & Plowright, R.K. (2020). Macroimmunology: The drivers and consequences of spatial patterns in wildlife immune defence. Journal of Animal Ecology, 89(4), 972–995. https://doi.org/10.1111/1365-2656.13166

Becker, D.J. & Hall, R.J. (2014). Too much of a good thing: resource provisioning alters infectious disease dynamics in wildlife. Biology Letters, 10(7), 20140309. https://doi.org/10.1098/rsbl.2014.0309

Briggs, C.J. & Hoopes, M.F. (2004). Stabilizing effects in spatial parasitoid–host and predator–prey models: a review. Theoretical Population Biology, 65(3), 299–315. https://doi.org/10.1016/j.tpb.2003.11.001

Brockhurst, M.A., Buckling, A. & Rainey, P.B. (2006). Spatial heterogeneity and the stability of host-parasite coexistence. Journal of Evolutionary Biology, 19(2), 374–379. https://doi.org/10.1111/j.1420-9101.2005.01026.x

Brooker, S., Clements, A.C. & Bundy, D.A. (2006). Global epidemiology, ecology and control of soil-transmitted helminth infections. Advances in Parasitology, 62, 221–261. https://doi.org/10.1016/S0065-308X(05)62007-6

Budischak, S.A., Hansen, C.B., Caudron, Q., Garnier, R., Kartzinel, T.R., Pelczer, I., Cressler, C.E., Van Leeuwen, A. & Graham, A.L. (2018). Feeding immunity: physiological and behavioral responses to infection and resource limitation. Frontiers in Immunology, 8(1914). https://doi.org/10.3389/fimmu.2017.01914

Cabaret, J., Gasnier, N. & Jacquiet, P. (1998). Faecal egg counts are representative of digestive-tract strongyle worm burdens in sheep and goats. Parasite, 5(2), 137–142. https://doi.org/10.1051/parasite/1998052137

Calenge, C. (2006). The package “adehabitat” for the R software: a tool for the analysis of space and habitat use by animals. Ecological Modelling, 197(3-4), 516–519. https://doi.org/10.1016/j.ecolmodel.2006.03.017

Carbayo, J., Martín, J. & Civantos, E. (2019). Habitat type influences parasite load in Algerian Psammodromus (Psammodromus algirus) lizards. Canadian Journal of Zoology, 97(2), 172–180. https://doi.org/10.1139/cjz-2018-0145

Clutton-Brock, T.H., & Pemberton, J.M. (2004). Individuals and populations.In: Clutton-Brock, T. H. & Pemberton, J. M. (eds.). Soay Sheep: Dynamics and Selection in an Island Population. Cambridge: Cambridge University Press, 1–16.

Clutton-Brock, T.H., Pemberton, J.M., Coulson, T., Stevenson, I.R. & MacColl, A.D.C. (2004). The sheep of St Kilda. In: Clutton-Brock, T. H. Cambridge University Press, 17–51.

Clutton-Brock, T.H., Price, O.F., Albon, S.D. & Jewell, P.A. (1992). Early development and population fluctuations in Soay sheep. Journal of Animal Ecology, 61(2), 381–396. https://doi.org/10.2307/5330

Costa, J.H.C., Costa, W.G., Weary, D.M., Machado Filho, L.C.P. & von Keyserlingk, M.A.G., (2016). Dairy heifers benefit from the presence of an experienced companion when learning how to graze. Journal of Dairy Science, 99(1), 562–568. https://doi.org/10.3168/jds.2015-9387

Coulson, G., Cripps, J.K., Garnick, S., Bristow, V. & Beveridge, I. (2018). Parasite insight: assessing fitness costs, infection risks and foraging benefits relating to gastrointestinal nematodes in wild mammalian herbivores. Philosophical Transactions of the Royal Society B: Biological Sciences, 373(20170197). https://doi.org/10.1098/rstb.2017.0197

Craig, B.H., Pilkington, J.G. & Pemberton, J.M. (2006). Gastrointestinal nematode species burdens and host mortality in a feral sheep population. Parasitology, 133(4), 485–496. https://doi.org/10.1017/S0031182006000618

Craig, B.H., Tempest, L.J., Pilkington, J.G. & Pemberton, J.M., (2008). Metazoan-protozoan parasite co-infections and host body weight in St Kilda Soay sheep. Parasitology, 135(4), 433–441. https://doi.org/10.1017/S0031182008004137

Crawley, M. J. (2017) The Flora of St Kilda. Hebridean Naturalist Supplement 1: 1-61. Curracag.

Crawley, M.J., Albon, S.D., Bazely, D.R., Milner, J.M., Pilkington, J.G. & Tuke, A.L. (2004). Vegetation and sheep population dynamics. In: Clutton-Brock, T. H. and Pemberton, J. M. (eds.). Soay Sheep: Dynamics and Selection in an Island Population. Cambridge University Press, 89–111.

Crawley, M.J., Pakeman, R.J., Albon, S.D., Pilkington, J.G., Stevenson, I.R., Morrissey, M.B., Jones, O.R., Allan, E., Bento, A.I., Hipperson, H., Asefa, G. & Pemberton, J. (2021). The dynamics of vegetation grazed by a food-limited population of Soay sheep on St Kilda. Journal of Ecology, 109(12), 3988–4006. https://doi.org/10.1111/1365-2745.13782

Cringoli, G., Rinaldi, L., Veneziano, V., Mezzino, L., Vercruysse, J. & Jackson, F. (2009). Evaluation of targeted selective treatments in sheep in Italy: Effects on faecal worm egg count and milk production in four case studies. Veterinary Parasitology, 164(1), 36–43. https://doi.org/10.1016/j.vetpar.2009.04.010

Daugherty, M.P. (2011). Host plant quality, spatial heterogeneity, and the stability of mite predator–prey dynamics. Experimental and Applied Acarology, 53(4), 311–322. https://doi.org/10.1007/s10493-010-9410-8

Díaz, S., Lavorel, S., de Bello, F., Quétier, F., Grigulis, K. & Robson, T.M. (2007). Incorporating plant functional diversity effects in ecosystem service assessments. Proceedings of the National Academy of Sciences, 104(52), 20684–20689. https://doi.org/10.1073/pnas.0704716104

Edwards, J. (1976). Learning to eat by following the mother in moose calves. American Midland Naturalist, 96(1), 229–232. https://doi.org/10.2307/2424583

Ezenwa, V.O. (2004). Interactions among host diet, nutritional status and gastrointestinal parasite infection in wild bovids. International Journal for Parasitology, 34(4), 535–542. https://doi.org/10.1016/j.ijpara.2003.11.012

Froy, H., Börger, L., Regan, C.E., Morris, A., Morris, S., Pilkington, J.G., Crawley, M.J., Clutton-Brock, T.H., Pemberton, J.M. & Nussey, D.H. (2018). Declining home range area predicts reduced late-life survival in two wild ungulate populations. Ecology Letters, 21(7), 1001–1009. https://doi.org/10.1111/ele.12965

Gardarin, A., Garnier, É., Carrère, P., Cruz, P., Andueza, D., Bonis, A., Colace, M.P., Dumont, B., Duru, M., Farruggia, A., Gaucherand, S., Grigulis, K., Kernéïs, É., Lavorel, S., Louault, F., Loucougaray, G., Mesléard, F., Yavercovski, N., & Kazakou, E. (2014). Plant trait–digestibility relationships across management and climate gradients in permanent grasslands. Journal of Applied Ecology, 51(5), 1207–1217. https://doi.org/10.1111/1365-2664.12293

Garnier, E., Cortez, J., Billès, G., Navas, M.L., Roumet, C., Debussche, M., Laurent, G., Blanchard, A., Aubry, D., Bellmann, A. & Neill, C. (2004). Plant functional markers capture ecosystem properties during secondary succession. Ecology, 85(9), 2630–2637. https://doi.org/10.1890/03-0799

Grime, J.P. (1998). Benefits of plant diversity to ecosystems: immediate, filter and founder effects. Journal of Ecology, 86(6), 902–910. https://doi.org/10.1046/j.1365-2745.1998.00306.x

Gulland, F.M.D. (1992). The role of nematode parasites in Soay sheep (Ovis aries L.) mortality during a population crash. Parasitology, 105(3), 493–503. https://doi.org/10.1017/S0031182000074679

Hayward, A.D., Wilson, A.J., Pilkington, J.G., Clutton-Brock, T.H., Pemberton, J.M. & Kruuk, L.E.B. (2011). Natural selection on a measure of parasite resistance varies across ages and environmental conditions in a wild mammal. Journal of Evolutionary Biology, 24(8), 1664–1676. https://doi.org/10.1111/j.1420-9101.2011.02300.x

Hayward, A.D., Wilson, A.J., Pilkington, J.G., Pemberton, J.M. & Kruuk, L.E. (2009). Ageing in a variable habitat: environmental stress affects senescence in parasite resistance in St Kilda Soay sheep. Proceedings of the Royal Society B: Biological Sciences, 276(1672), 3477–3485. https://doi.org/10.1098/rspb.2009.0906

Hill, M.O., Mountford, J.O., Roy, D.B. & Bunce, R.G.H. (1999). Ellenberg’s indicator values for British plants. ECOFACT, Vol 2 Technical Annex. Huntingdon, Institute of Terrestrial Ecology, 46 pp. https://nora.nerc.ac.uk/id/eprint/6411/

Hutchings, M. R., Milner, J. M., Gordon, I. J., Kyriazakis, I. & Jackson, F. (2002). Grazing decisions of Soay sheep, Ois aries, on St Kilda: a consequence of parasite distribution? Oikos, 96(2), 235–244. https://doi.org/10.1034/j.1600-0706.2002.960205.x

Hutchings, M.R., Gordon, I.J., Kyriazakis, I. & Jackson, F. (2001). Sheep avoidance of faeces-contaminated patches leads to a trade-off between intake rate of forage and parasitism in subsequent foraging decisions. Animal Behaviour, 62(5), 955–964. https://doi.org/10.1006/anbe.2001.1837

Illius, A.W., Albon, S.D., Pemberton, J.M., Gordon, I.J. & Clutton-Brock, T.H. (1995). Selection for foraging efficiency during a population crash in Soay sheep. Journal of Animal Ecology, 64(4), 481–492. https://doi.org/10.2307/5651

Jewell, P.A., Milner, C. & Morton Boyd, J. (eds.) (1974). Island Survivors: The Ecology of the Soay Sheep of St. Kilda. Cambridge University Press, Cambridge.

Johnson, P.T., Wood, C.L., Joseph, M.B., Preston, D.L., Haas, S.E. & Springer, Y.P. (2016). Habitat heterogeneity drives the host-diversity-begets-parasite-diversity relationship: evidence from experimental and field studies. Ecology Letters, 19(7), 752–761. https://doi.org/10.1111/ele.12609

Jones, O.R., Crawley, M.J., Pilkington, J.G. & Pemberton, J.M. (2005). Predictors of early survival in Soay sheep: cohort-, maternal-and individual-level variation. Proceedings of the Royal Society B: Biological Sciences, 272(1581), 2619–2625. https://doi.org/10.1098/rspb.2005.3267

Jones, O.R., Pilkington, J.G. & Crawley, M.J. (2006). Distribution of a naturally fluctuating ungulate population among heterogeneous plant communities: ideal and free? Journal of Animal Ecology, 75(6), 1387–1392. https://doi.org/10.1111/j.1365-2656.2006.01163.x

Kleyer, M., Bekker, R.M., Knevel, I.C., Bakker, J.P., Thompson, K., Sonnenschein, M., Poschlod, P., Van Groenendael, J.M., Klimeš, L., Klimešová, J. Klotz, S., Rusch, G.M., Hermy, M., Adriaens, D., Boedeltje, G., Bossuyt, B., Dannemann, A., Endels, P., Götzenberger, L. et al. (2008). The LEDA Traitbase: a database of life-history traits of the Northwest European flora. Journal of Ecology, 96(6), 1266–1274. https://doi.org/10.1111/j.1365-2745.2008.01430.x

Laliberté, E., & Legendre, P. (2010) A distance-based framework for measuring functional diversity from multiple traits. Ecology, 91, 299–305. https://doi.org/10.1890/08-2244.1

Laliberté, E., Legendre, P., & Shipley, B. (2014). FD: measuring functional diversity from multiple traits, and other tools for functional ecology. R package version 1.7-19.

Lindgren, F., Rue, H. & Lindström, J. (2011). An explicit link between Gaussian fields and Gaussian Markov random fields: the stochastic partial differential equation approach. Journal of the Royal Statistical Society: Series B (Statistical Methodology), 73(4), 423–498. https://doi.org/10.1111/j.1467-9868.2011.00777.x

Lynsdale, C.L., Mumby, H.S., Hayward, A.D., Mar, K.U. & Lummaa, V. (2017). Parasite-associated mortality in a long-lived mammal: Variation with host age, sex, and reproduction. Ecology and Evolution, 7(24), 10904–10915. https://doi.org/10.1002/ece3.3559

McGill, B.J., Enquist, B.J., Weiher, E. & Westoby, M. (2006). Rebuilding community ecology from functional traits. Trends in Ecology & Evolution, 21(4), 178–185. https://doi.org/10.1016/j.tree.2006.02.002

Milner, J.M., Albon, S.D., Illius, A.W., Pemberton, J.M. & Clutton-Brock, T.H. (1999). Repeated selection of morphometric traits in the Soay sheep on St Kilda. Journal of Animal Ecology, 68(3), 472–488. https://doi.org/10.1046/j.1365-2656.1999.00299.x

Mitsunaga, T. & Fujii, K. (1997). The effects of spatial and temporal environmental heterogeneities on persistence in a laboratory experimental community. Researches on Population Ecology, 39(2), 249–260. https://doi.org/10.1007/BF02765271

Morand, S., Krasnov, B.R. & Poulin, R. (eds.) (2007). Micromammals and Macroparasites: From Evolutionary Ecology to Management. Springer. https://doi.org/10.1007/978-4-431-36025-4

Murdoch, W.W. (1977). Stabilizing effects of spatial heterogeneity in predator-prey systems. Theoretical Population Biology, 11(2), 252–273. https://doi.org/10.1016/0040-5809(77)90028-4

Nielsen, M.K., Kaplan, R.M., Thamsborg, S.M., Monrad, J. & Olsen, S.N. (2007). Climatic influences on development and survival of free-living stages of equine strongyles: implications for worm control strategies and managing anthelmintic resistance. The Veterinary Journal, 174(1), 23–32. https://doi.org/10.1016/j.tvjl.2006.05.009

Nussey, D.H., Watt, K., Pilkington, J.G., Zamoyska, R. & McNeilly, T.N. (2012). Age-related variation in immunity in a wild mammal population. Aging Cell, 11(1), 178–180. https://doi.org/10.1111/j.1474-9726.2011.00771.x

Pakeman, R.J. (2014). Leaf dry matter content predicts herbivore productivity, but its functional diversity is positively related to resilience in grasslands. PloS One, 9(7), p.e101876. https://doi.org/10.1371/journal.pone.0101876

Powell, R. (2000). Animal home ranges and territories and home range estimators. In: Boitani, L. & Fuller, T. K. (eds.). Research Techniques in Animal Ecology: Controversies and Consequences. Columbia University Press, 65–110.

Putfarken, D., Dengler, J., Lehmann, S. & Härdtle, W. (2008). Site use of grazing cattle and sheep in a large-scale pasture landscape: A GPS/GIS assessment. Applied Animal Behaviour Science, 111(1-2), 54–67. https://doi.org/10.1016/j.applanim.2007.05.012

R Core Team. (2021). R: A language and environment for statistical computing. R Foundation for Statistical Computing, Vienna, Austria. https://www.R-project.org/.

Regan, C.E., Pilkington, J.G., Pemberton, J.M. & Crawley, M.J. (2016). Sex differences in relationships between habitat use and reproductive performance in Soay sheep (Ovis aries). Ecology Letters, 19(2), 171–179. https://doi.org/10.1111/ele.12550

Roeber, F., Jex, A.R. & Gasser, R.B. (2013). Advances in the diagnosis of key gastrointestinal nematode infections of livestock, with an emphasis on small ruminants. Biotechnology Advances, 31(8), 1135–1152. https://doi.org/10.1016/j.biotechadv.2013.01.008

Roscher, C., Schumacher, J., Gubsch, M., Lipowsky, A., Weigelt, A., Buchmann, N., Schmid, B. & Schulze, E.D. (2012). Using plant functional traits to explain diversity–productivity relationships. PloS one, 7(5), p.e36760. https://doi.org/10.1371/journal.pone.0036760

Rue, H., Martino, S. & Chopin, N. (2009). Approximate Bayesian inference for latent Gaussian models by using integrated nested Laplace approximations. Journal of the Royal Statistical Society: Series B (Statistical Methodology), 71(2), 319–392. https://doi.org/10.1111/j.1467-9868.2008.00700.x

Smith, L.A., White, P.C., Marion, G. & Hutchings, M.R. (2009). Livestock grazing behavior and inter-versus intraspecific disease risk via the fecal–oral route. Behavioral Ecology, 20(2), 426–432. https://doi.org/10.1093/beheco/arn143

Sparks, A.M., Watt, K., Sinclair, R., Pilkington, J.G., Pemberton, J.M., Johnston, S.E., McNeilly, T.N. & Nussey, D.H. (2018). Natural selection on antihelminth antibodies in a wild mammal population. The American Naturalist, 192(6), 745–760. https://doi.org/10.1086/700115

Sweeny, A.R., Albery, G.F., Venkatesan, S., Fenton, A. & Pedersen, A.B. (2021). Spatiotemporal variation in drivers of parasitism in a wild wood mouse population. Functional Ecology, 35(6), 1277–1287. https://doi.org/10.1111/1365-2435.13786

Sweeny, A.R., Clerc, M., Pontifes, P.A., Venkatesan, S., Babayan, S.A. & Pedersen, A.B. (2021). Supplemented nutrition decreases helminth burden and increases drug efficacy in a natural host– helminth system. Proceedings of the Royal Society B, 288(1943), 20202722. https://doi.org/10.1098/rspb.2020.2722

Sweeny, A.R., Corripio-Miyar, Y., Bal, X., Hayward, A.D., Pilkington, J.G., McNeilly, T.N., Nussey, D.H. & Kenyon, F. (2022). Longitudinal dynamics of co-infecting gastrointestinal parasites in a wild sheep population. Parasitology, 149(5), 593–604. https://doi.org/10.1017/S0031182021001980

Tilman, D. (2001). Functional diversity. Encyclopedia of Biodiversity, 3(1), 109–120. https://doi.org/10.1016/B978-0-12-384719-5.00061-7

Verheyden, H., Richomme, C., Sevila, J., Merlet, J., Lourtet, B., Chaval, Y. & Hoste, H. (2020). Relationship between the excretion of eggs of parasitic helminths in roe deer and local livestock density. Journal of Helminthology, 94, e159. https://doi.org/10.1017/S0022149X20000449

Vicari, M., Puentes, A., Granath, G., Georgeff, J., Strathdee, F. & Bazely, D.R. (2018). Unpacking multi-trophic herbivore-grass-endophyte interactions: feedbacks across different scales in vegetation responses to Soay sheep herbivory. The Science of Nature, 105(66). https://doi.org/10.1007/s00114-018-1590-9

Vlassoff, A., Leathwick, D.M. & Heath, A.C.G. (2001). The epidemiology of nematode infections of sheep. New Zealand Veterinary Journal, 49(6), 213–221. https://doi.org/10.1080/00480169.2001.36235

Weiher, E., Van Der Werf, A., Thompson, K., Roderick, M., Garnier, E. & Eriksson, O. (1999). Challenging Theophrastus: a common core list of plant traits for functional ecology. Journal of Vegetation Science, 10(5), 609–620. https://doi.org/10.2307/3237076

Weiss, K.C. & Ray, C.A. (2019). Unifying functional trait approaches to understand the assemblage of ecological communities: synthesizing taxonomic divides. Ecography, 42(12), 2012–2020. https://doi.org/10.1111/ecog.04387

Williams, B. & Warren, J. (2004). Effects of spatial distribution on the decomposition of sheep faeces in different vegetation types. Agriculture, Ecosystems & Environment, 103(1), 237–243. https://doi.org/10.1016/j.agee.2003.09.016

Wilson, K., Grenfell, B.T., Pilkington, J.G., Boyd, H.E.G. & Gulland, F.M.D. (2004). Parasites and their impact. In: Clutton-Brock, T. H. & Pemberton, J. M. (eds.). Soay Sheep: Dynamics and Selection in an Island Population. Cambridge University Press, 113–165.

Wilson, P.J., Thompson, K. & Hodgson, J.G. (1999). Specific leaf area and leaf dry matter content as alternative predictors of plant strategies. The New Phytologist, 143(1), 155–162. https://doi.org/10.1046/j.1469-8137.1999.00427.x

Zouyed, I., Cabaret, J. & Bentounsi, B. (2016). Climate influences assemblages of abomasal nematodes of sheep on steppe pastures in the east of Algeria. Journal of Helminthology, 92(1), 34–41. https://doi.org/10.1017/S0022149X16000845

