## Supplementary Figure 1 for "Age specific impacts of vegetation functional traits on gastro-intestinal nematode parasite burdens in a large herbivore"

***Supplementary Figure S1.*** *Vegetation rastas of each vegetation functional trait showing spatial variation in values within the study area. Vegetation rastas were generated from the community weighted means of each grid square of the study area. Each grid square is 100 m^2^. For all traits, higher values are green and lower values are light pink. Eastings and northings are standardized coordinates (in units of 1 m) from the southern- and western-most points of the study area.*

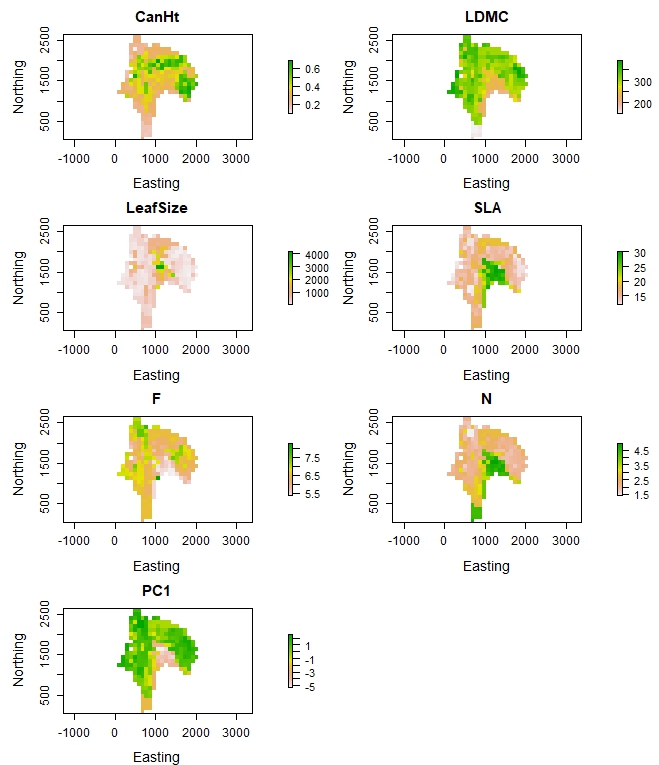
