## Supplementary Figure 2 for "Age specific impacts of vegetation functional traits on gastro-intestinal nematode parasite burdens in a large herbivore"

***Supplementary Figure S2. Confidence intervals (95%) for models of (a) adults and (b) yearlings****. Base model includes sex, and weight as fixed effects, and year as a random effect. Base models for adults also include individual as a random effect and age as a continuous fixed effect. The similarity in confidence intervals of each term across models suggests that the effects are consistent following inclusion of random effects. Each model, and the combination of terms included therein, is assigned a colour and, as not all models include all terms, not all models can be compared for a given term. Traits are insignificant in the model if the confidence interval crosses the central dashed line. Positive estimates indicate a positive effect of the term on FEC.*

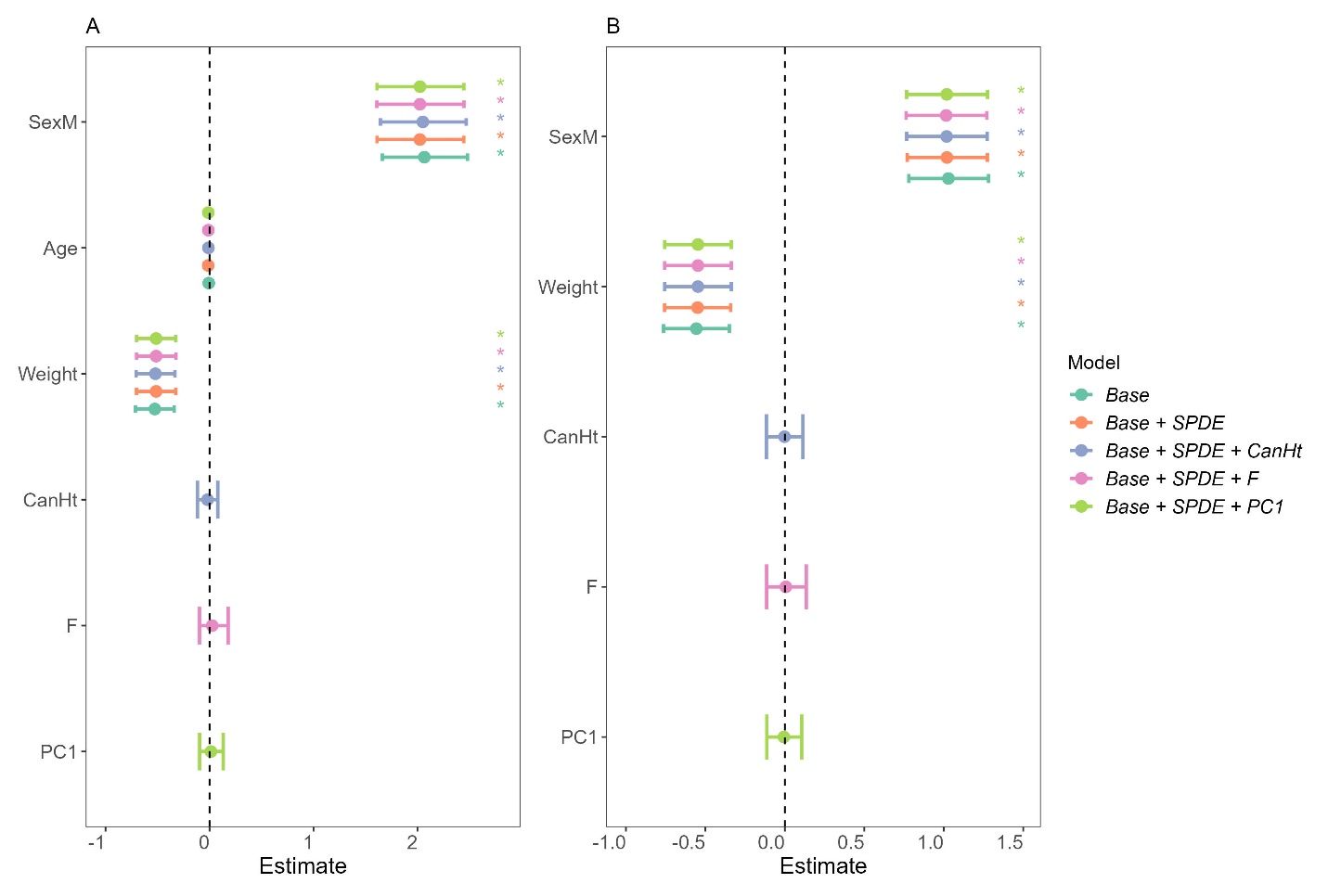
