## Supplementary Table 1 for "Age specific impacts of vegetation functional traits on gastro-intestinal nematode parasite burdens in a large herbivore"

***Supplementary Table S1.*** *Correlation matrix of the vegetation variables used.*

|  | **CanHt** | **LDMC** | **LeafSize** | **SLA** | **F** | **N** |
| --- | --- | --- | --- | --- | --- | --- |
| **CanHt** | 1 | - | - | - | - | - |
| **LDMC** | 0.18156 | 1 | - | - | - | - |
| **LeafSize** | -0.075734 | -0.33605 | 1 | - | - | - |
| **SLA** | -0.25039 | -0.63020 | 0.56965 | 1 | - | - |
| **F** | -0.15115 | 0.54239 | -0.30515 | -0.56025 | 1 | - |
| **N** | 0.37592 | -0.84074 | 0.51618 | -0.82990 | -0.66138 | 1 |
