## Supplementary Table 2 for "Age specific impacts of vegetation functional traits on gastro-intestinal nematode parasite burdens in a large herbivore"

***Supplementary Table S2****. Summary of PCA with a) loadings from PCA and b) contributions of each variable to dimensions in PCA.*

a) Loadings from PCA

|  | **PC1** | **PC2** | **PC3** | **PC4** | **PC5** | **PC6** |
| --- | --- | --- | --- | --- | --- | --- |
| **CanHt** | 0.15107 | -0.87277 | 0.10067 | -0.29668 | 0.25308 | 0.23077 |
| **LDMC** | 0.45192 | 0.012363 | 0.35674 | 0.71506 | 0.12689 | 0.37543 |
| **LeafSize** | -0.34070 | -0.046988 | 0.86318 | -0.10560 | -0.34214 | -0.091766 |
| **SLA** | -0.48179 | 0.039663 | 0.13561 | 0.20654 | 0.81108 | -0.21773 |
| **F** | 0.38846 | 0.47284 | 0.28175 | -0.58837 | 0.37552 | 0.24280 |
| **N** | -0.52331 | 0.103797 | -0.14055 | -0.026296 | -0.062573 | 0.83127 |

b) Contributions from PCA

|  | **PC1** | **PC2** | **PC3** | **PC4** | **PC5** | **PC6** |
| --- | --- | --- | --- | --- | --- | --- |
| **CanHt** | 2.2822 | 76.172 | 1.0135 | 8.8021 | 6.4049 | 5.3253 |
| **LDMC** | 20.423 | 0.015285 | 12.726 | 51.130 | 1.6102 | 14.095 |
| **LeafSize** | 11.608 | 0.22079 | 74.508 | 1.1151 | 11.706 | 0.84209 |
| **SLA** | 23.213 | 0.15731 | 1.8390 | 4.2657 | 65.786 | 4.7408 |
| **F** | 15.090 | 22.357 | 7.9383 | 34.617 | 14.102 | 5.8951 |
| **N** | 27.385 | 1.0774 | 1.9754 | 0.069147 | 0.39154 | 69.102 |
